# Tox_(R)CNN: Deep Learning-Based Nuclei Profiling tool For Drug Toxicity Screening

**DOI:** 10.1101/334557

**Authors:** Daniel Jimenez-Carretero, Vahid Abrishami, Laura Fernández-de-Manuel, Irene Palacios, Antonio Quílez-Álvarez, Alberto Díez-Sánchez, Miguel Angel del Pozo, María C. Montoya

## Abstract

Toxicity is an important factor in failed drug development, and its efficient identification and prediction is a major challenge in drug discovery. We have explored the potential of microscopy images of fluorescently labeled nuclei for the prediction of toxicity based on nucleus pattern recognition. Deep learning algorithms obtain abstract representations of images through an automated process, allowing them to efficiently classify complex patterns, and have become the state-of-the art in machine learning for computer vision. Here, deep convolutional neural networks (CNN) were trained to predict toxicity from images of DAPI-stained cells pre-treated with a set of drugs with differing toxicity mechanisms. Different cropping strategies were used for training CNN models, the nuclei-cropping-based Tox-CNN model outperformed other models classifying cells according to health status. Tox-CNN allowed automated extraction of feature maps that clustered compounds according to mechanism of action. Moreover, fully automated region-based CNNs (RCNN) were implemented to detect and classify nuclei, providing per-cell toxicity prediction from raw screening images. We validated both Tox-(R)CNN models for detection of pre-lethal toxicity from nuclei images, which proved to be more sensitive and have broader specificity than established toxicity readouts. These models predicted toxicity of drugs with mechanisms of action other than those they had been trained for and were successfully transferred to other cell assays. The Tox-(R)CNN models thus provide robust, sensitive, and cost-effective tools for *in vitro* screening of drug-induced toxicity. These models can be adopted for compound prioritization in drug screening campaigns, and could thereby increase the efficiency of drug discovery.

**Author summary:** Visualization of nuclei using different microscopic approaches has for decades allowed the identification of cells undergoing cell death, based on changes in morphology, nuclear density, etc. However, this human-based visual analysis has not been traslated into quantitative tools able to objectively measure cytotoxicity in drug-exposed cells. We asked ourselves if it would be possible to train machines to detect cytotoxicity from microscopy images of fluorescently stained nuclei, without using specific toxicity labeling. Deep learning is the most powerful supervised machine learning methodology available, with exceptional abilities to solve computer vision tasks, and was thus selected for the development of a toxicity quantification tool. Two convolutional neural networks (CNN) were developed to classify cells based on health status: Tox-CNN, relying on prior cell segmentation and cropping of nuclei images, and Tox-RCNN which carries out fully-automated cell detection and classification. Both Tox-(R)CNN classification outputs provided sensitive screening readouts that detected pre-lethal toxicity and were validated for a broad array of toxicity pathways and cell assays. Tox-(R)CNN approaches excel in affordability and applicability to other in vitro toxicity readouts and constitute a robust screening tool for drug discovery.

## Introduction

Toxicity is a major cause of failure in drug development and causes costly withdrawals of drugs from the market. Drug development productivity would be greatly improved if cytotoxic compounds were identified during early *in vitro* screening [1–4]. Drug-induced cytotoxic effects lead to changes in cell and nuclear morphology which are characteristic of the specific cell-death pathway involved, the best characterized being apoptosis, necrosis, and autophagy [5–7]. The field has advanced with the establishment of high content screening (HCS) techniques and the emergence of toxicity reporters revealing specific biochemical pathways triggered during cell death programs or measuring metabolic cell function [8–10]. However, toxicity reporters are often limited to assess the specific biochemical pathways for which they were designed [11–13], and they are thus unlikely to capture the wide variety of toxic effects that can be triggered by different drugs in screening campaigns. Toxicity-screening approaches have combined multi-parametric image analysis of fluorescently labeled nuclei with the use of toxicity reporters in advanced machine learning pipelines [14–18]. However, toxicity reporters increase experimental complexity, thus reducing throughput and increasing screening costs. There is therefore an urgent need to develop broad specificity, cost-effective *in vitro* toxicity assays for incorporation in the primary screening phases of drug development. Cytotoxic effects have classically been visually identified from cell and nuclear morphology [5–7]. However, the complexity and variability of toxicity-associated morphological patterns has so far hindered their systematic and quantitative analysis and thus prevented their use as standalone toxicity screening endpoints. Although nuclear fluorescence staining forms the basis of most high content cell-based assays, its use is normally limited to image segmentation and nuclei counting to score cell-loss due to lethal toxicity [19,20], thus disregarding the wealth of information contained in images of fluorescently labeled nuclei. In an effort to exploit this information for the quantification of pre-lethal toxicity, we have explored state-of the-art machine learning tools for automated pattern recognition. The success of classical learning-based computer vision methods relies heavily on extraction and selection of a reasonable set of relevant features that are highly discriminative of the phenotypes being studied. Feature selection requires in-depth knowledge of the phenotype under study, which is hindered in the current application by the complexity and great variety of drug-induced toxicity-associated nuclei organization patterns. The most recent major advance in machine learning is deep convolutional neural networks (CNN) which, similar to a brain, have multiple layers of interconnected artificial neurons [21,22]. Through an automated process, deep neural networks learn abstract representations of raw images from pixel information as a progressive hierarchy of sub-images, from which they extract features that can be used to classify complex patterns in a supervised manner. CNNs can thus automate the critical steps of feature extraction and selection by learning to extract high-level features based on spatial relationships, which has enabled them to outperform other machine learning methods in computer vision tasks, as demonstrated for several challenging biomedical applications [23–28]. Thus, deep technology seemed well suited to the analysis and prediction of drug toxicity in images of fluorescently labeled nuclei.

Here, we present novel deep-learning approaches for in vitro cell-based toxicity assessment. The Tox-(R)CNN approaches proved to efficiently predict a broad spectrum of toxicity mechanisms from different drugs and cell lines. The main strength of these tools is their unique ability to predict toxicity based exclusively on nuclei staining; this offers the advantage of improved affordability and applicability of toxicity prediction. The attraction of the Tox-(R)CNN tools relies on their high potential to enable sensitive and efficient compound prioritization based on detection of pre-lethal toxicity in primary screening campaigns.

## Results

### CNN can predict cell toxicity based on images of fluorescently stained nuclei

To test if CNNs can predict cell toxicity based exclusively on nuclear staining, we designed an experimental assay in which HL1 cells were treated with reference compounds at different concentrations. We used previously established dose-curves to guarantee varying degrees of toxicity outcome. The included reference compounds cover a range of cytotoxic effects: the DNA targeting genotoxic drugs cyclophosphamide (Ciclo), 5-fluorouracil (5Fluo), and doxorubicin (Doxo); the apoptosis-triggering drug staurosporine (Staur); the enzyme inhibitors acetaminophen (Aceta) and sunitinib (Sunit), which are not associated with a specific toxicity mechanism; the uncoupler of mitochondrial oxidative metabolism FCCP; and the microtubule stabilizer Taxol, which inhibits mitosis. Cells were labeled with the DNA-specific fluorescent probe DAPI and imaged with an automated confocal microscope, revealing a great variety of nuclear patterns induced by the different compounds (Fig 1A). The standard toxicity readout of nucleus count (Num Nuc) revealed cell loss due to drug-induced lethal effects, but did not reveal the great variety of toxic effects associated with the reference compounds assayed (Fig 1B). As reference standards, we analyzed caspase 3/7 nuclear translocation and Mitotracker cytoplasmic intensity (Fig 1C,D), both of which evidenced dose-dependent toxic effects promoted by the compounds tested. To assess toxicity independently of cell density and to enable detection of pre-lethal toxicity, we implemented a deep CNN architecture for the estimation of cell health status from microscopy images of DAPI-stained nuclei (Fig 1E). Since we are aiming at a cell-based toxicity assessment, we used standard image analysis procedures to segment nuclei and cytoplasm according to the DAPI signal and used the segmentation to crop images (see Materials and Methods). Based on the regions of interest included in the resulting image crop, four image cropping strategies were designed and used to train independent CNN models: nuclei (Nuc), nuclei and cytoplasm (Cell), nuclei and 3 adjacent pixels (Nuc_Ring), and nuclei, cytoplasm, and background (All) (Fig 1F). The resulting image crops, each containing a mass-centered cell, were used as inputs for CNN-based toxicity prediction. For each cell, the CNN output layer delivers a “health status” score, which determines a binary classification: *healthy* or *toxicity affected*. As a supervised learning technique, the CNN required images labeled according to the expected output (*healthy* or *toxicity affected*) as ground-truth images for training. However, the reference standards analyzed for this purpose, caspase 3/7 and Mitotracker, did not qualify as general toxicity labels because neither efficiently captured all the drug-induced cytotoxic effects included in our experimental assay: both yielded poor resolution of Taxol-induced cytotoxicity and of the kinetic effects produced by Doxo and Aceta. As an alternative strategy, we produced a training dataset by labeling image crops according to the treatment exposure; cells from untreated wells were labeled *healthy*, whereas those from wells treated with the highest drug concentrations were labeled *toxicity affected* (S1A Fig). The percentage of cells classified as *healthy* by the different CNN models served as the per-well measurement of “general” toxicity (Fig 1G-K). To allow comparative evaluation, Z-scores were obtained from the percentage of cells classified as *healthy* by the CNN model for each compound-treated well by normalizing to the negative control (DMSO). Z-scores, displayed so that positive values depict toxic effects, revealed that all the readouts obtained with CNN models were more efficient at predicting toxicity than the classical Num Nuc readout (Fig 1G), with CNN_Nuc displaying the highest Z-scores for the drugs Doxo, Staur, and Taxol. In spite of the high degree of uncertainty introduced into the impure training set, the CNN *healthy* scores efficiently revealed dose-response toxicity curves for all the drugs tested in the assay (Fig 1H-K), providing better resolution than those obtained from Num Nuc or the caspase or Mitotracker readouts (Fig 1B-D). At high toxicant doses, CNN models trained with information from nuclei only (CNN_Nuc) underwent an unexpected drop in toxicity prediction. This was not observed in CNN models trained with cytoplasmic information (CNN_Cell/_Nuc_Ring/_All), which performed better at these doses due to their enhanced ability to reveal DNA release into the cytoplasm during necrosis induced by high drug concentrations. However, the CNN_Nuc model proved to be the best readout for early prediction of toxicity, since it outperformed other cropping strategies at sub-lethal drug concentrations, where there is no reduction in Num Nuc indicating significant cell loss. Consistent with this pattern, plotting treatments with above-threshold Z-scores as “toxic hits” revealed CNN_Nuc to be the most sensitive method for detecting early toxicity yielding 100 toxic hits out of 184 treated wells (Fig 2A), which outperformed all other CNN models. In turn, CNN_Cell _, Nuc_Ring, and _ALL readouts yielded 87, 86, and 85 hits, respectively (Fig 2B,C and S1B Fig). The standard toxicity readouts Num nuc, caspase and Mitotracker detected 27, 81, and 6 toxic hits, respectively (Fig 2D-F). All CNN models yielded significant Z-scores for Taxol at 0.1μM, demonstrating broader application than established readouts; caspase detected toxicity effects only at 2μM Taxol, whereas Mitotracker and Num Nuc did not detect significant Taxol toxicity. The higher sensitivity of CNN Nuc compared with the other CNN models was further evidenced by the lower half-maximal toxic concentration values (EC50) obtained for the mild toxicant 5Fluo (1.47μM and 2.73μM for CNN_Nuc and CNN_Cell, respectively) (Fig 2). Based on these results, nuclear crops were used as inputs for CNN models in all subsequent studies, and from here on CNN refers to these CNN_Nuc models.

**Fig 1.**
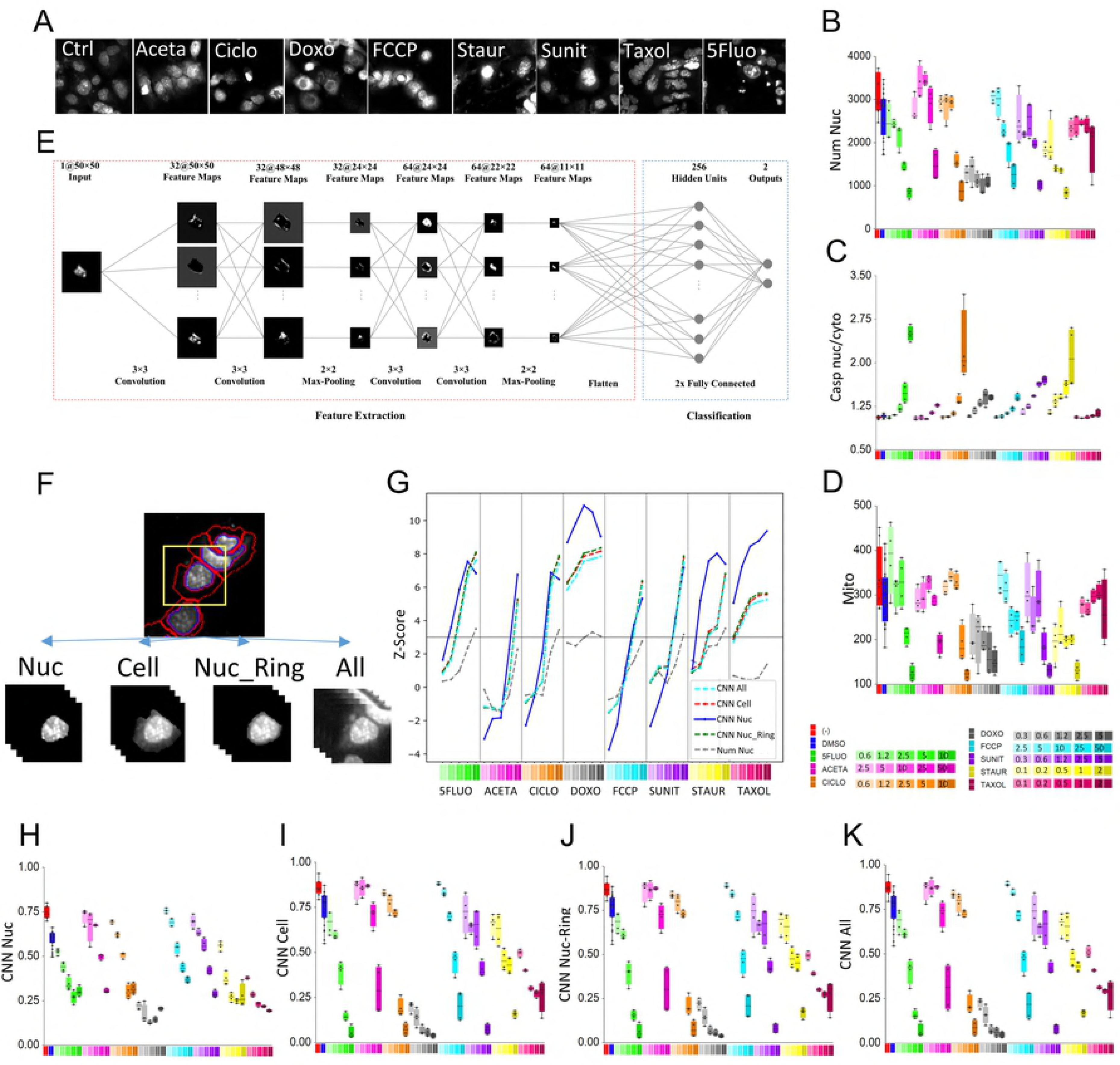
Toxicity prediction using deep CNN strategies compared with established readouts. HL1 cells treated or not (-) with the indicated concentrations of compounds (µM) or DMSO were processed as described in the Materials and Methods. (A) Representative fluorescence microscopy images of DAPI-stained cells treated or not (Ctrl) with the highest concentrations of the indicated compounds in a reference experiment used for CNN training. (B-D) Boxplots of per-well toxicity assessments from established measurements: nucleus count (Num Nuc) (B), caspase 3/7 nucleus:cytoplasm ratio (Casp nuc/cyto) (C), and Mitotracker cytoplasmic intensity (Mito) (D). (E) CNN architecture for predicting health status from single-cell image crops, as described in Materials and Methods. (F) Cropping strategies; representative image crops are shown of nucleus (Nuc), nucleus+cytoplasm (Cell), nucleus+margin (Nuc_Ring), and nucleus+cytoplasm+background+neighboring cells (All). (G) Plot displaying mean toxicity readouts of replicate wells, obtained from the percentage of *healthy* cells predicted by the different CNN models (CNN_Nuc, _Nuc_Ring, _Cell, _All) and the standard nuclei counting (Num Nuc) for the different treatments indicated. For each well, toxicity readouts were obtained by computing Z-scores (normalizing to DMSO-treated wells) with adjustment of the sign to display toxic effects as positive values. (H-K) Boxplots of per-well toxicity assessments from CNN-based predictions: percentage of cells classified as *healthy* by the CNN_Nuc (H), CNN_Cell (I), CNN_Nuc_Ring (J), and CNN_All (K) models.

**Fig 2.**
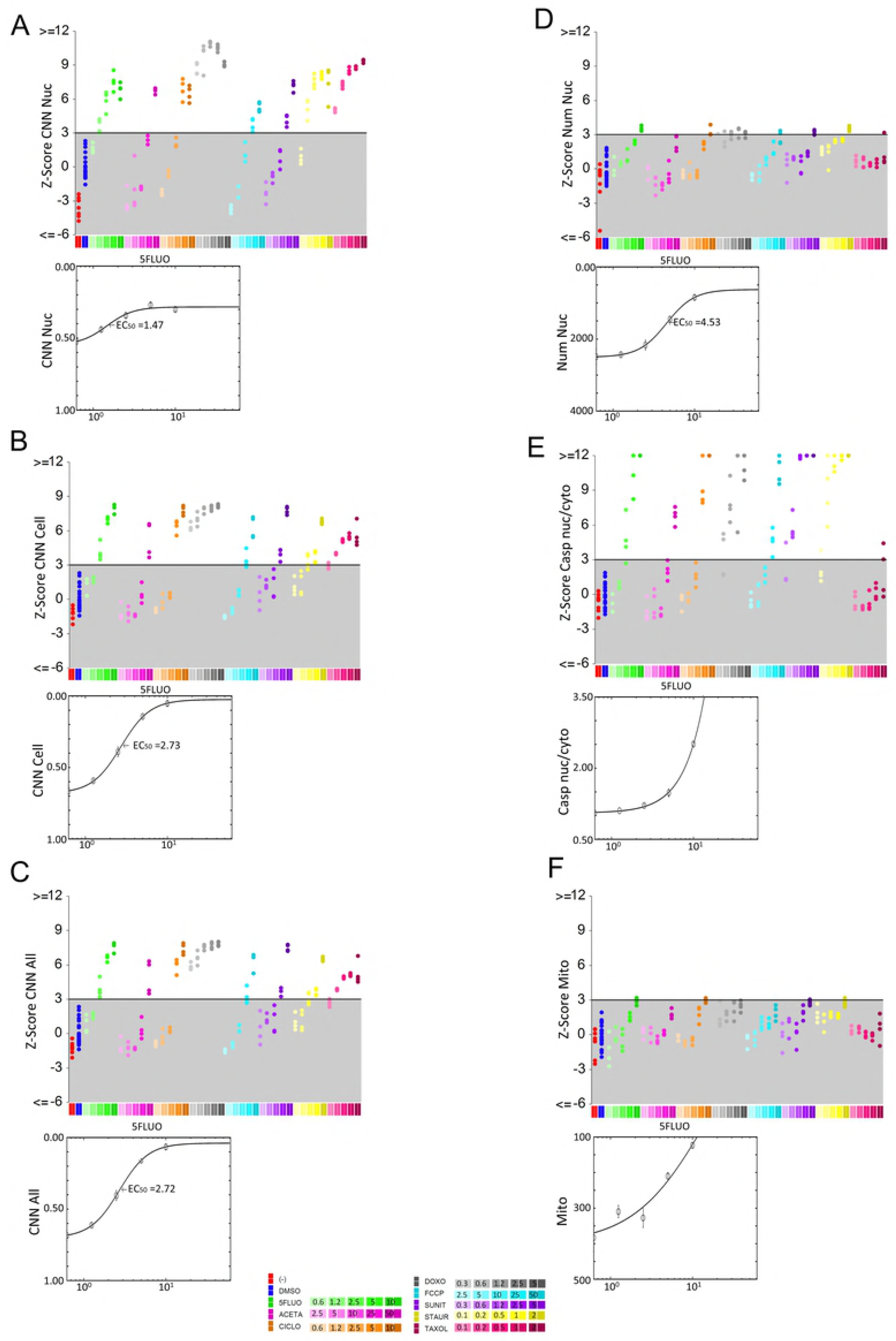
Sensitivity of CNN-based toxicity prediction compared with standard toxicity readouts. HL1 cells treated or not (-) with DMSO or the indicated concentrations of drugs (µM) were processed as described in the Materials and Methods. Plots display individual well toxicity readouts (top) and the 5-Fluorouracil dose-response curve fitted from well-averages, including the EC_50_ (bottom). (A-C) CNN-based toxicity readouts: CNN_Nuc (A), CNN_Cell (B), and CNN_All (C). (D-F) Standard toxicity readouts: Nuclei counting by standard image segmentation (Num Nuc) (D), mean Caspase 3/7 nucleus:cytoplasm ratio (Casp Nuc/Cyto) (E), and mean Mitotracker cytoplasmic intensity (Mito) (F). For each well, toxicity readouts were obtained by computing Z-scores (normalizing to DMSO-treated wells) with adjustment of the sign to display toxic effects as positive values. Z-scores > 3 represent toxic hits.

### Region-based CNN (RCNN) for fully automated toxicity prediction

To avoid relying on external image segmentation and cropping procedures while providing a per-cell toxicity prediction, we undertook an alternative cell-based deep-learning approach incorporating automated object detection. An RCNN was implemented for the automated localization and classification of individual nuclei using raw images as input, instead of crops. The framework, based on the Faster RCNN algorithm [29], includes a region proposal network (RPN) that uses features extracted from the last convolutional layer of a CNN to detect bounding boxes around individual candidate cell, which are then classified as *healthy, toxicity-affected*, or *background* (Fig 3A,B). We trained the RCNN model with cell bounding box coordinates obtained with the standard segmentation procedure used in the CNN approach (see Materials and Methods). A set of 7 independent experiments in HL1 cells treated with the eight reference drugs, including two experiments in which untreated cells were cultured at different confluencies, was used for in-parallel training of the RCNN and CNN mixed models for toxicity prediction using balanced datasets of *healthy* and *toxicity affected* labeled nuclei (Tox-CNN and Tox-RCNN_balanced). Resulting CNN predictions showed that the Tox_RCNN_balanced model, unlike Tox_CNN, erroneously predicted toxicity of untreated cells grown at low densities (Fig 3C). To prevent the models from learning to predict toxicity as a reduction in cell number due to drug-induced lethality, we trained an additional mixed model that included extra images of untreated cells cultured at different densities (120 extra healthy training wells), hereafter referred to as Tox_RCNN. The unbalanced and balanced Tox-RCNN models (Num Nuc RCNN and Num Nuc RCNN_balanced) detected a similar number of objects (nuclei); moreover, the number was consistent with the number obtained by the standard segmentation procedure (Num Nuc) at regular cell densities (Fig 3C-E), demonstrating the efficiency of automated detection by RCNN. Tox_RCNN slightly overestimated cell number at low confluencies, probably due to the recognition of cellular debris that are discarded by the image processing procedure. In an independent experiment, both Tox-CNN and Tox-RCNN mixed models successfully classified untreated cells at very low densities as healthy and efficiently predicted the toxic effects of drugs and high DMSO concentrations (Fig 3C), further demonstrating their independence from cell-density fluctuations. Tox-(R)CNN models performed efficiently in the test wells from an experiment used for training (Fig 3D) and in one independent experiment with HL1 cells at higher confluency (S2A Fig). Overall, Tox-(R)CNN models were consistently more sensitive than Num Nuc at predicting drug toxicity, with Tox-CNN outperforming Tox-RCNN classification in most cases. Even though these models were trained in HL1 cells, they suitably predicted toxicity from these drugs in two other cell lines, EAHY926 (Fig 3E) and MEVEC (S2B Fig), thus confirming the applicability of Tox-(R)CNN models for the prediction of toxicity in different cell types. Together, these results demonstrate the robustness of deep-learning-based toxicity prediction with regard to inter-experimental and intra-experimental variability, thus confirming Tox-(R)CNN as powerful screening tools.

**Fig 3.**
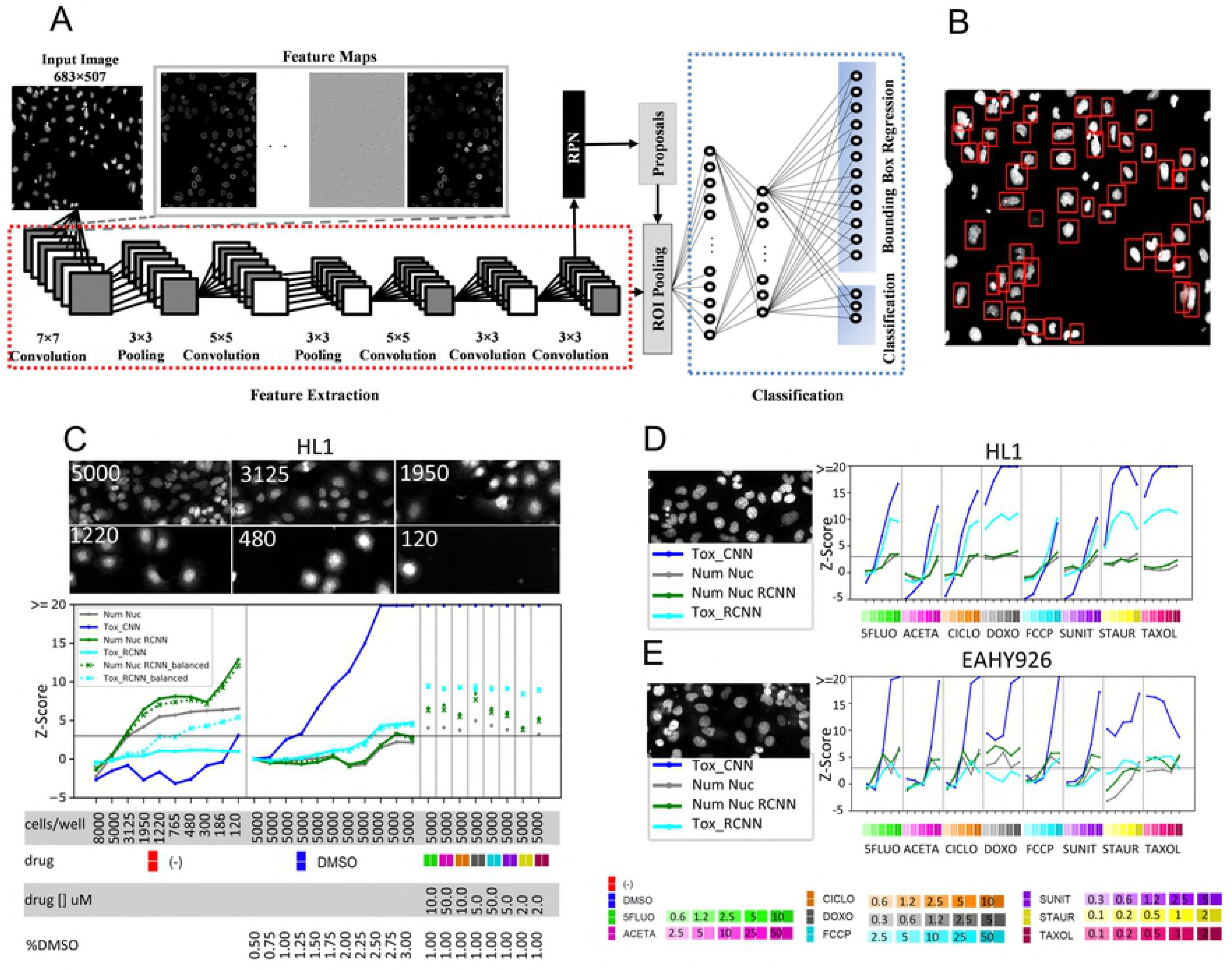
Region-based CNN implementation and evaluation of (R)CNN deep-learning toxicity-assessment approaches. (A) RCNN architecture for automated detection of cells and prediction of their health status from micrographs of DAPI fluorescence, as described in the Materials and Methods. (B) Example of nuclei bounding boxes resulting from the region proposal network included in the RCNN framework. (C) HL1 cells were seeded at the indicated densities (cells/well). (D,E) HL1 and EAHY926 cells were seeded at 5000 cells/well. (C-E) 24h after seeding, cells were treated or not (-) with the indicated concentrations of DMSO (%) or the indicated drugs (µM) and processed as described in the Materials and Methods. Representative images are shown of untreated cells at the indicated cell-seeding densities (cells/well). Plots display mean toxicity readouts of four replicate wells, obtained from the percentage of *healthy* cells predicted by the CNN_Nuc (Tox_CNN) and RCNN (Tox_RCNN_balanced and Tox_RCNN) mixed models, and from nuclei counting by standard image segmentation (Num Nuc), or by using RCNN-based automated detection (Num Nuc RCNN) from Tox_RCNN training. For each well, toxicity readouts were obtained by computing Z-scores (normalizing to DMSO-treated wells) with adjustment of the sign to display toxic effects as positive values.

### Validation of Tox-(R)CNN models for toxicity prediction of different drugs and mechanisms of action

To demonstrate the value of these Tox-(R)CNN models as tools for broad toxicity prediction, we performed a new screening in which HL1 cells were treated with a panel of 24 drugs, including those used to train the CNN model and additional drugs acting through several mechanisms. Tox-(R)CNN mixed models sensitively predicted the outcome of toxic compound-treatments, thus proving their ability to reveal the toxicity of compounds for which they have not been trained (Fig 4A). Conveniently, Tox-CNN enabled the automated extraction of features from pixel intensity values, which were used for unsupervised hierarchical clustering of compounds (see Materials and Methods). Tox-CNN features clustered compounds in a biologically meaningful manner, even though the models were not trained for this purpose (Fig 4B). Statins (lovastatin and simvastatin) clustered together, as did the ionophores FCCP and monesin, which produce ROS and mitochondrial toxicity. The DNA synthesis inhibitors 5Fluo, gemcitabine, and mitomicine also group in the same cluster. Apoptotic death inducers (staurosporine, thapsigargin, bortezomib, and imatinib) were clustered closely together with other drugs of unknown mechanism. The DNA intercalating anthracyclines epirubicin and doxorubicin also clustered together. These findings confirm not only that deep CNN are able to perceive general toxic effects, but also that their ability to learn feature representations provides useful knowledge for exploring toxicity mechanisms.

**Fig 4.**
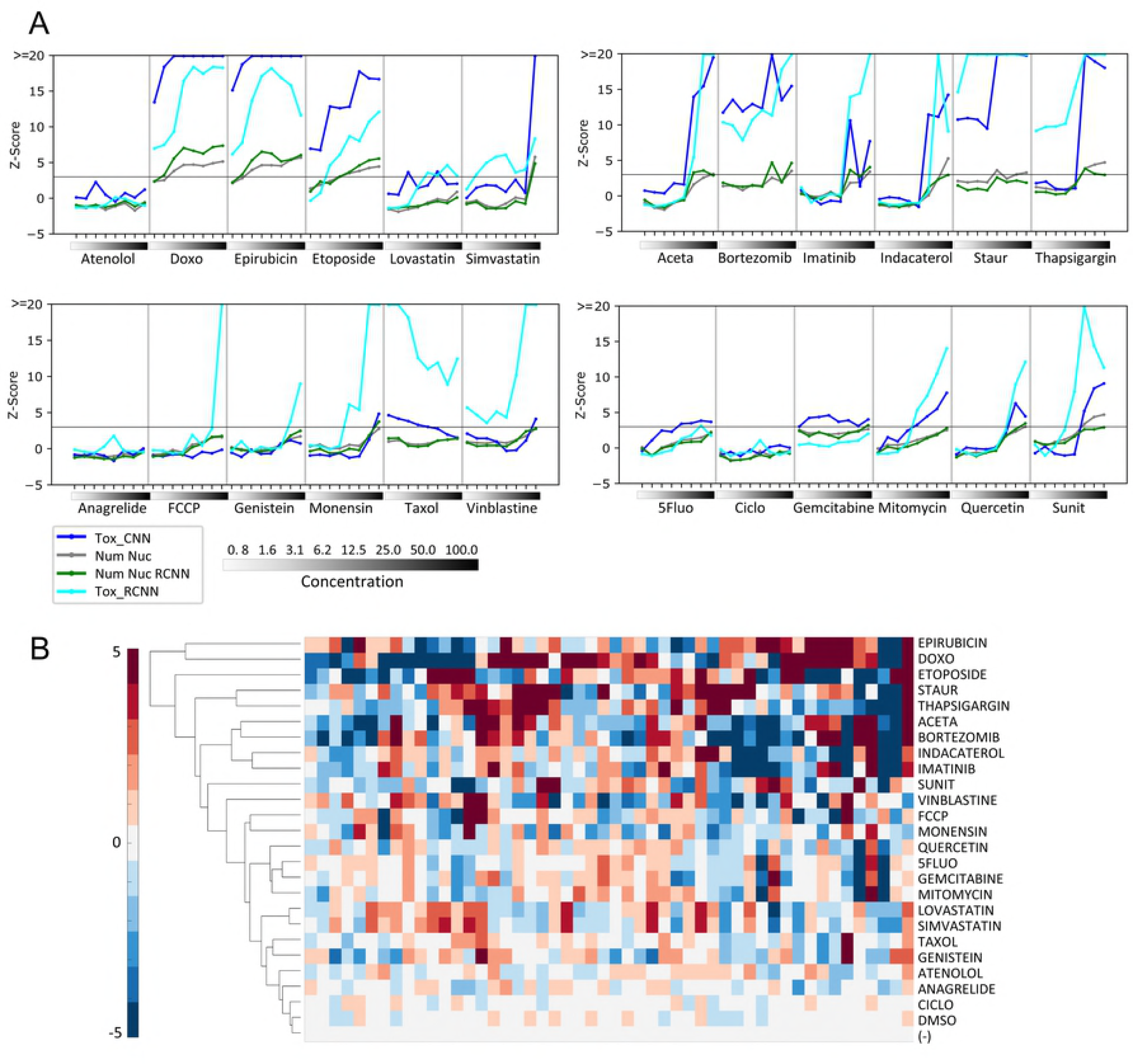
Deep (R)CNN model validation to predict toxicity and to extract knowledge for clustering of drugs based on their toxicity mechanisms. HL1 cells treated with one of 24 compounds or DMSO at the concentrations indicated (µM) were processed as described in the Materials and Methods. (A) Plots displaying mean toxicity readouts of four replicate wells, obtained from percentage of *healthy* cells predicted by the CNN_Nuc (Tox_CNN) or RCNN (Tox_RCNN) mixed models, and from nuclei counting by standard image segmentation (Num Nuc), or by using RCNN-based automated detection (Num Nuc RCNN). For each well, toxicity readouts were obtained by computing Z-scores (normalizing to DMSO-treated wells) with adjustment of the sign to display toxic effects as positive values. (B) Hierarchical clustering of features obtained with the Tox_CNN model from HL1 cells treated with 25µM of the indicated drugs or 0.78µM Taxol; untreated (-); DMSO, control.

### Validation of Tox-(R)CNN models as toxicity screening tools

To further evaluate the Tox-(R)CNN deep-learning models as screening tools for prioritizing compounds based on their toxicity potential, we re-analyzed a pre-accomplished HCS of primary pancreatic cancer associated fibroblasts (pan-CAFs). Among several assay-specific labels, most of which are irrelevant here (not shown), this HCS included DAPI staining in the assay for both image segmentation and nuclei counting as a toxicity endpoint. A transfer learning strategy was applied to the Tox-(R)CNN models delivering (Tr_Tox_(R)CNN) models. Training strategy was designed to allow the use of pre-run screens lacking reference toxicity-inducing treatments (see Materials and Methods). In brief, the training dataset was produced from images from drug-treated wells with a significantly reduced cell number, which were labeled *toxicity affected*, while cells from DMSO treated wells were labeled *healthy,* since no untreated cells were available for training. Interestingly, compounds #19 and #33 (anagrelide and quercetin) were negligibly lethal according to nuclei counting, but were predicted by Tox-CNN to be toxic at high concentrations (Fig 5A,B), demonstrating the greater sensitivity of deep-learning-based prediction compared with Num Nuc. Toxicity of these compounds was confirmed by re-testing in primary cardiac fibroblasts (S3 Fig). Even though Tr_Tox_CNN performed better that the Tox-CNN model, the latter was still a more sensitive predictor of toxic effects than nuclei counting, demonstrating the value of these tools. Nuclei counting by both the transferred and original Tox_RCNN models (Tr_Tox_RCNN and Tox RCNN) was consistent with the standard procedure (Num Nuc), revealing that object detection was performing adequately. However, the original Tox-RCNN model was a poor toxicity predictor in pan-CAFs compared with the transferred RCNN model (Tr_Tox_RCNN), further evidencing the need for a transfer-learning approach for RCNN toxicity predictions in cell lines different from those used for training. The screening comprised 60 drugs at 8 concentrations (480 treatments) and yielded 102 toxicity hits (mean Z score > 3) based on nuclei counting (Fig 5C). In contrast Tr_Tox_CNN and Tr_Tox_RCNN identified 127 and 126 toxic wells, respectively (Fig 5D,E). Z-scores computed for the transferred Tr_(R)CNN models were plotted independently for all compounds screened (S4 Fig). These results further demonstrate the superior performance and sensitivity of deep Tox-(R)CNN toxicity predictions over classical screening endpoints based on nuclei staining.

**Fig 5.**
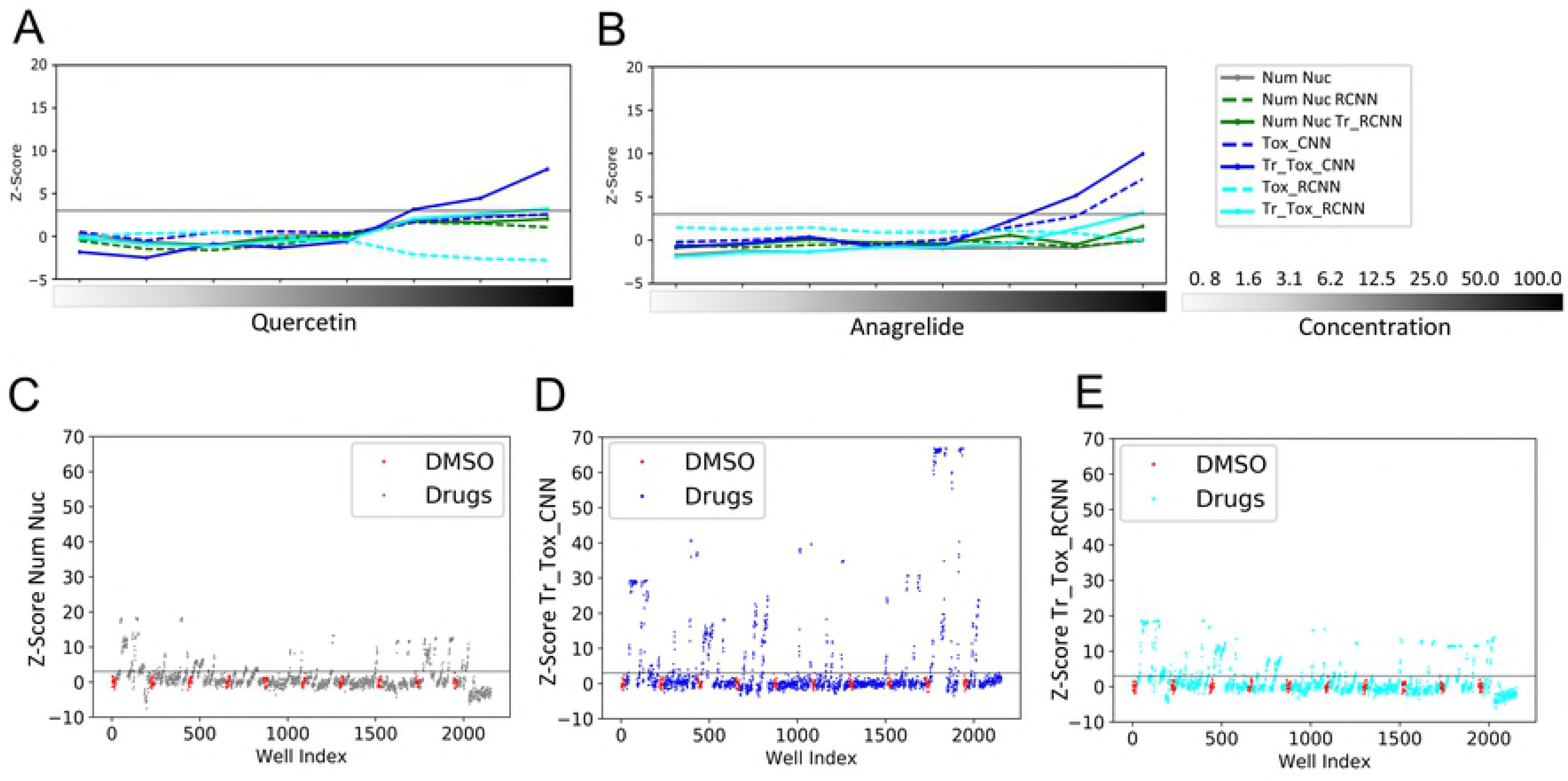
Validation of deep (R)CNN models for general drug toxicity screening. Pancreatic CAFs treated with 60 compounds or DMSO at the concentrations indicated (µM) were processed as described in the Materials and Methods. (A-B) Plots display mean toxicity readouts of replicate wells for quercetin (#19, A) and anagrelide (#33, B), obtained from the percentage of *healthy* cells predicted by the CNN_Nuc (Tox_CNN) or RCNN (Tox_RCNN) mixed models before (dashed lines) and after (solid lines) transfer learning, and from nuclei counting by standard image segmentation (Num Nuc), or by using RCNN-based automated detection (Num Nuc RCNN) before (dashed lines) and after (solid lines) transfer learning. (C-E) Plots showing individual well toxicity readouts obtained after transfer learning from: nuclei counting using standard image segmentation (Num Nuc) (C), percentage of *healthy* cells predicted by CNN (Tr_Tox_CNN) (D), and percentage of *healthy* cells predicted by RCNN (Tr_Tox_RCNN) (E). Negative control, DMSO-treated wells, are highlighted in red. For each well, toxicity readouts were obtained by computing Z-scores (normalizing to DMSO-treated wells) with adjustment of the sign to display toxic effects as positive values.

## Discussion

There is a need to incorporate highly predictive toxicity assays into primary in vitro high-throughput screening in order to reduce the attrition of drug candidates at later phases of the drug discovery pipeline. In vitro cytotoxicity assessment is normally limited to measuring the number of viable cells per well. However, single-cell readouts make better outputs that avoid sources of experimental error such as non-homogeneities in cell-dispensing, drug-induced proliferative effects, and heterogeneous responses of different cell sub-populations, which could be misinterpreted if only well averages are examined. The toxicity research field has therefore been directed towards finding novel cell labels and readouts that distinguish between different cytotoxicity mechanisms [8–13]. Nevertheless, the use of toxicity reporters has not gained broad acceptance because it adds experimental complexity, thus reducing throughput and increasing screening costs. Here, we have established tools that predict cell toxicity based on the analysis of fluorescently labeled nuclei. These tools outperformed outputs that rely on toxicity reporters or cell counting. Since nuclei staining is common to most high content cell based assays, this tool has broad applicability for toxicity prediction in HCS, even in pre-accomplished screens, as demonstrated here.

Over recent years, deep learning approaches have been successfully deployed in computer vision tasks and constitute the state-of-the-art tools for supervised machine learning. Several key features give deep learning approaches an advantage over other machine learning methodologies for toxicity analysis. First, by learning to represent data with multiple levels of abstraction in an unsupervised manner (i.e. without human-based programming) it avoids cumbersome knowledge-based feature engineering. This makes deep learning approaches independent of prior in depth knowledge of the target phenotypes, and therefore more suitable for broad toxicity prediction of multiple drugs with differing mechanisms by HCS. Second, their ability to learn intricate patterns improves recognition, feature extraction, and classification from noisy images, providing accuracy. This has led to deep technology outperforming other machine learning methods and human-based analysis, as demonstrated for several challenging biomedical applications [23–28]. Third, using transfer learning methods, networks can be updated to classify new datasets with limited data training [26,27,30], and thus makes them suitable for predicting toxicity in pre-accomplished screenings. Accordingly, the deep-learning approaches presented here successfully predict toxicity of a broad spectrum of toxicity-associated patterns in HCS images of fluorescently labeled nuclei, proving to be more sensitive than established toxicity reporters and readouts for the recognition of pre-lethal toxicity. The Tox-(R)CNN models suitably predicted toxicity in cell lines and for compounds different from the ones used for training. Moreover, for very different cell lines, pre-trained Tox_(R)CNN models can be subject to a simple transfer-learning approach that does not require toxicity controls, increasing the performance of toxicity predictions; this was successfully achieved here in the pre-accomplished screening with primary pancreatic cancer associated fibroblasts (panCAF). Although CNN models were more sensitive and transferable and performed better, the use of RCNN models is justified by their independence from external segmentation and image cropping. We also demonstrate the utility of CNNs for extracting knowledge (as feature maps) that allows comparative analysis of the toxicity mechanisms exerted by different drugs in a screen. Previous cell-based toxicity screening approaches have combined multi-parametric image analysis of fluorescently labeled nuclei with the use of toxicity reporters in advanced machine learning pipelines [14–18]. The main strength of the tools presented here is their unique ability to predict toxicity based exclusively on nuclei staining, which offers the advantage of improving affordability and applicability of toxicity prediction. HCS is now an established tool for phenotypic drug discovery; in this setting, the deep-learning approaches presented here will promote a better use of HCS technology for toxicity assessment. The relevance of the deep learning approaches presented here relies on their high potential to enable sensitive and efficient compound prioritization based on detection of pre-lethal toxicity. They thus provide affordable cytotoxicity counter-screens for high throughput primary campaigns and could allow academic screening centers and pharma companies to discard cytotoxic compounds during primary screening and hit-to-lead drug development campaigns, thereby increasing the efficiency of drug discovery.

## Materials and Methods

### Cell culture and reagents

Mouse cardiac muscle HL-1 cells purchased from Merck Millipore were grown on fibronectin(25µg/ml)/gelatin(1mg/ml) coated dishes with 10% fetal bobine serum (FBS) in Claycomb medium (Sigma-Aldrich). Mouse embryonic ventricular endocardial cells (MEVEC) were kindly provided by Dr. de la Pompa [31] and cultured on 0.1% gelatin coated flasks in 10% FBS supplemented DMEM. EAHY926 cells were kindly provided by Dr. Edgell and maintained in 10% FBS supplemented DMEM. PanCAF were obtained from Dr. Hidalgo and cultured in RPMI with 20% FBS. Primary pig cardiac fibroblasts were isolated from fresh surgical samples by collagenase tissue digestion as described in [32]. Fibroblasts were grown to 80% confluency in flasks containing supplemented DMEM. All culture media were supplemented with 10% FBS (except PanCAFs, which had 20% FBS), 100 U/ml penicillin, 100 μg/ml streptomycin and 2 mM L-glutamine and refreshed every 2-3 days. Mycoplasma tests were performed bimonthly for all cell line cultures.

Fluorescence staining reagents including DAPI (4’, 6-diamidine-2-fenilindol), Mitotracker Orange and Cell Event 3/7 caspase Green for detection of nuclei, active respiratory cell mitochondria and apoptosis respectively, were purchased from Invitrogen. Dimethyl sulfoxide vehicle (DMSO); Acetaminophen (ACETA); Doxorubicin (DOXO); carbonylcyanide p-trifluoromethoxyphenylhydrazone (FCCP); Sunitinib; Staurosporine (STAUR); Paclitaxel (TAXOL); Imatinib; Thapsigargin; Gemcitabine; Quercetin; Atenolol; Simvastatin; Genistein; Vinblastine; Monensin; Anagrelide; Epirubicin; Etoposide and Lovastatin were from Sigma. Ciclophosphamide (CICLO), 5-Fluorouracil (5FLUO)and Sunitinib malate (SUNIT) were from Tocris. Indacaterol and Bortezomib were kinldy provided by Dr. Blanco, Experimental Therapeutics Programme at CNIO.

### Assay procedure

Cells were seeded on 384-well plates (5000 cells/well, otherwise specified), after 24h compounds were added to wells at 8 serial concentrations in 4 replicate wells using layout shown in S1A Fig. To avoid evaporation-related edge effects external rows were filled with PBS and not used for the assay. Compounds were dissolved in DMSO (final concentration of 1% across the entire assay, otherwise specified). Cells were maintained in culture with compounds for 24h, and then stained for Caspase 3/7 and/or MitoTracker prior fixation with 4% PFA for 10 min at RT and final DAPI staining. Imaging of DAPI stained nuclei and fluorescent toxicity probes was performed with an Opera automated confocal microscope (Perkin Elmer) fitted with an NA=0.7 water immersion objective at a magnification of 20×;.

### Image Processing

Image processing algorithm was developed using Definiens Developer version XD2.4 (Definiens AG, Germany). Nuclei and cytoplasmic regions were first segmented based on differential contrast of DAPI intensity. Cells with a nuclear size bellow 0.3 times or over twice the mean of nuclear sizes per field were considered as debris and reclassified as non-cellular regions. An ID was univocally assigned to each cell the number of cells per well was computed. Toxicity readouts based on fluorescent reporters were extracted as Caspase 3/7 nucleus/cytoplasm intensity ratio (Casp nuc/cyto), and Mitotracker mean intensity in the cytoplasmic region (Mito), where specified.

Images from DAPI stained nuclei were cropped and saved individually assuring one cropped image per single mass-centered cell, thus conserving univocal cell ID. Four different strategies of cropping images were established (Fig 1F):

a. Nucleus (Nuc): Pixels from nuclear region preserve their intensity, while pixels from other regions including cytoplasm, neighbour cells or background are set to zero. Unless specified, this cropping strategy is used for training Tox_CNN models and classification purposes throughout the manuscript.
b. Nucleus and ring (Nuc_Ring): Pixels from nuclear region including a thin margin of 3 pixels containing a small region of the cell cytoplasm adjacent to the nuclei preserve their intensity. Pixels from other regions including the remaining cytoplasm, neighbour cells or background are set to zero. This cropping strategy was designed to include properties of nuclear border that could benefit the classification task.
c. Cell (Cell): Pixels from nuclear and cytoplasmic regions preserve their intensity. Pixels from other regions, including neighbour cells or background are set to zero. This cropping strategy incorporates information of DAPI signal beyond the nuclear boundaries.
d. All regions (All): All pixels within the crop preserve their intensity, which include those from the cell nucleus and cytoplasm, the background and eventually the fractions of neighboring cells.

The four strategies were set to a fixed size of 50×50 pixel crops, thus guaranteeing a proper inclusion of complete nuclei based on nuclear sizes and image pixel sizes. To avoid off-centered nuclei crops, those nuclei with a distance to field image boundaries of less than 50 pixels were excluded from the cropping extraction and further analysis. Aditionally, bounding box coordinates from segmented cells were also extracted for training automatic object detection by RCNN.

### Convolutional neural network (Tox-CNN)

We designed the toxicity convolutional neural network with 4 convolutional and 2 fully connected layers (Fig 1E) to classify single-cell images as “healthy” or “toxicity affected”. The Rectified Linear Unit (ReLU) activation function is applied between each layer except the output dense layer, which uses a softmax activation function to provide a separate probability for each of the classes. Convolutional layers convolve a 3×3 kernel over some input to produce 32, 32, 64, and 64 feature maps, respectively. To reduce the number of features and the computational complexity of the network, we introduced two max-pooling layers with a window size of 2×2 after convolutional layers 2 and 4. Additionally, to avoid overfitting, we included two dropout blocks after convolutional layers 2 and 4, and another one next to the first fully connected layer. Dropout deactivates some neurons randomly with a probability of 25% during the weight update cycle. The final max-pooling layer is then flattened and followed by two densely connected layers with 512 and 2 features. Finally, we applied a softmax activation function to the output of last fully connected layer to calculate the probability for each class label. The total number of parameters to learn is equal to 4,031,458, most of them belong to the first fully connected layer. We used ADADELTA algorithm [33] to adjust the learning rate automatically. To increase the number of data and avoid overfitting, we augmented images by applying random rotations in the range of [0°, 20°], horizontal shifts in the range of [0, 0.2 ×; *Image*_*width*_], vertical shifts in the range of [0, 0.2 ×;*Image*_*height*_], and horizontal/vertical flips, where *Image*_*width*_ and *Image*_*height*_ show width and height of input images, respectively. By default, the modifications were applied randomly, so not every image will be changed every time. Images were normalized to zero mean and unit variance before feeding them into the network.

### Region based Convolutional Neural Networks (Tox-RCNN)

We used state-of-the-art Faster Region-Based CNN [29] (RCNN) to both detect and classify cells as “healthy” or “toxicity affected” from entire images. Faster RCNN is composed of two modules: a Regional Proposal Network (RPN) and a RCNN network. RPN is a Fully Convolutional Network (FCN) [34] which proposes square regions within an image that may contain objects of interest without considering their classes while RCNN network classifies the object proposals from RPN into one of the classes (or background), and refines the bounding boxes’ coordinates of the final proposals. We used the original Faster RCNN architecture [29] without any significant modifications except for the number of outputs for the classification and regression layers since we have two classes. Therefore, the classification layer has 18 (9×2) outputs, and the regression layer has 36 (9×4) outputs (coordinates).

### Label-free toxicity annotation of images

To train the network for single-cell toxicity prediction, a set of cells need to be labelled as “healthy” and “toxicity affected”. The uncertainty of the toxicity state of individual cells due to lack of bona-fide toxicity reporters hamper the possibility of creating a pure and clean training dataset. Therefore, cells were labeled according to the known or expected toxic response of different extreme treatments; a set of untreated cells (or cells under harmless treatment) were labelled as “healthy”, and a set of cells treated with known toxic compounds at the highest concentrations were labelled as “toxicity affected”. This labelling strategy minimize the amount of manual supervision needed to perform the tedious task of creating a large annotated dataset for training, avoiding also the use of any toxicity labels such as fluorescent reporters that always provide partial information since they are unable to detect all the toxic effects. These “healthy” and “toxicity affected” sets are initially conformed by all cells in a selection of wells correspondent to the appropriate extreme treatments. Both sets were balanced, where indicated by removing all cells from randomly selected entire images. The final output is a field-based treatment-driven training dataset that represents two groups/classes with the (expected) highest mean difference in terms of toxicity state. A training-strategy avoiding overfitting was undertaken to properly deal with minimized, yet still present mislabeling of cells affected by toxicity in conditions with harmless or no treatment, as well as resilient cells in extreme harmful conditions.

### Training Tox-(R)CNN

Training of both Tox-CNN and Tox-RCNN models need the outputs obtained from the image processing routine developed in Definiens (see Image processing section). CNN approach used cropped single cell 50 ×; 50 pixel images; and RCNN approach used full fields (683 ×; 507 pixel images) and bounding-box coordinates of cells. The initial experiment pursuing the comparison of CNN performance from 4 cropping strategies (Nuc, Nuc_Ring, Cell and All) used images from one experimental plate, where four correspondent trainings were performed in parallel. The creation of central Tox-(R)CNN mixed models involved images from 7 HL1 experiments including several conditions: untreated cells, cells treated with DMSO, and cells treated with up to 8 drugs with known toxic effects at different concentrations. All drugs used demonstrated toxic effects in previous experiments where dose curves were fixed. Three of these experiments were designed in a way that allows the classifier to learn that healthy cells can also grow at low densities. Training dataset was created as detailed in the previous section, labelling single cells from wells without any treatment as “healthy”, and cells from treated wells with highest concentrations of the 8 available compounds as “toxicity affected”. In each plate, only half of these wells per condition were used for training, and the rest were bound to test. In total, Tox-CNN mixed model was trained using 739,727 cells (image crops) covering 8 compounds (“toxicity affected”) and untreated cells (“healthy”). Bounding-box coordinates of training cells, together with the correspondent 7,487 and 10,883 entire images (fields) were used to train the balanced and final Tox-RCNN mixed models, respectively. We set the number of epochs to 120 and the batch size to 256 when training the Tox-CNN, and used 750,000 iterations for Tox-RCNN.

### Transfer learning Tr_Tox-(R)CNN

For CAF screening, we used deep transfer learning [35] to adapt Tox-(R)CNN mixed models to a different cell line improving toxicity prediction. With this strategy, an existing pre-trained network is fine-tuned, avoiding training an entire neural network from scratch and reusing low level feature-detectors already learned. Therefore, we froze the weights of the first two convolutional layers in both (R)CNN approaches and retrained the rest of the layers to adapt to this new dataset. For this screening, since there is no prior information about the dose-response curve and expected toxicity, we used a different strategy to create the training set, but again following the guidelines detailed before (see Label-free toxicity annotation of images). First, we selected drugs with at least one concentration with a significant toxic effect scored as a significant reduction of the number of cells per well (Z-score>3). Then, for each selected drug, cells treated with the two highest concentrations of that drug were included in the training set, and labelled as “toxicity affected”. Since untreated wells are not included in regular screenings, cells from half of the DMSO-treated wells were included in the “healthy” training set, which it corresponds to the harmless condition in this specific screening. This resulted in a training set with 170,775 instances (6,057 field-images). We followed this general strategy for generating training sets, to ensure that it can be used in any assay where no prior information about toxic effects of compounds included in the screening is available. We used 25 epochs to re-train the Tox-CNN model, and 45,000 iterations to update the weights of the Tox-RCNN model; conforming the transferred models for toxicity prediction in PanCAF screening (Tr_Tox-(R)CNN models).

### Tox-(R)CNN classification and evaluation

We evaluated the performance of Tox-(R)CNN models on several independent experiments (test sets). Cells out of the training set coming from experiments that were partially used for training were also employed for testing purposes: treatments with intermediate concentration of drugs were never included in training set, and not all wells from extreme treatments were selected for training. Tox-CNN model classified crop images at the input as “healthy” or “toxicity affected” based on the probabilistic scores obtained at the output for both classes. Tox-RCNN model return object detections which were classified as “healthy”, “toxicity affected” or “background” (considered as non-cell detections and discarded from further analyses) in entire field images.

### Toxicity readouts

Well-based toxicity measurements were constructed from Tox-(R)CNN predictions by computing the percentage of cells classified as “healthy” in each well. Standard toxicity measurements obtained from fluorescent reporters (Caspase 3/7 and Mitotracker) were aggregated in well-based values by computing the mean. Nuclei count obtained by image processing and RCNN-derived counting of nuclear detections were also reported for each well. For each type of toxicity measure (*x*), we computed Z-scores by subtracting the mean (*μ*) and then dividing by the standard deviation (*σ*) of negative control (DMSO treated wells):

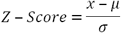

Finally, we adjust the sign of the outputs to get increasing values for toxic effects, thus obtaining final well-based toxicity readouts that allow direct comparisons. Values with a sign adjusted Z-score over 3 were considered as hits [36].

### Hierarchical clustering

Tox-CNN features from the first fully connected layer (512 features/sample) were extracted for all cells in 4-plate screening of HL-1 cells with 24 drugs. Features obtained from cells that were either untreated, DMSO-treated, or treated with a non-extreme toxic concentration of the drugs (25 µM) (except for Taxol in which a 0,781 µM concentration was used) were analyzed by PCA to select the 50 most informative features for further processing. Mean well-based features were later normalized (Z-score with respect to untreated condition) and aggregated in order to obtain mean values per treatment/condition. Finally, these feature vectors were assembled by hierarchical clustering (Euclidean distance metric and weighted average linkage) to generate the hierarchical tree. Clusters along both dimensions (features and treatment/condition) were displayed in a heatmap of feature vector values including dendrograms representing the multilevel hierarchy obtained.

### EC50

Half maximal effective concentration (EC50) is the concentration of a drug which induces a response halfway between the observed baseline and maximum effect of that drug. For the present work, the drug responses used for the EC50 calculations are the different toxicity effects, as indicated. EC50 were computed by fitting a Hill Equation sigmoid curve to the dose-response values and estimating the correspondent EC50 Hill Equation parameter [37]. Dose-response values and adjusted curves are displayed in Fig 2 and S1B Fig in such way that visual increasing values depict toxic effects for all toxicity measurements in order to allow proper comparison. Response axes are fixed to [1; 0] for CNN predictions, [4,000; 0] for nuclei count, [0.5; 3.5] for the Caspase ratio and [500; 100] for Mitotracker measurements. EC50 values were not calculated in cases where standard sigmoidal curve fitting was inappropriate within the existing range of drug doses (Fig 2E,F).

### Software and hardware resources

We used Keras [38] on the Theano backend [39] to develop the Tox-CNN model presented here. To construct the Tox-RCNN model we used a python implementation of Faster RCNN [29] which is developed on Caffe [40]. All experiments were run on a standard PC with an Intel Xeon CPU E5-2643 @ 3.30 GHz, 32 GB working memory, and using a 16 GB NVIDIA Quadro K4000 GPU to speed up the computations. Training time for the Tox-CNN model was 11 hours; transfer learning took 2 hours. Training time for the Tox-RCNN mixed model and transfer learning were 187 hours and 9 hours, respectively. Boxplots and well-based dot plots were created with NCSS Statistical Software (version 11), dose-response curve adjustments and EC50 calculations were computed using MATLAB (R2017a), and mean Z-score plots were depicted with Python.

### Code and data availability

Source code, documentation, trained models, and exemplary datasets are available via Github; Tox-CNN: https://github.com/nielintos/Tox-CNN, Tox-RCNN: https://github.com/nielintos/Tox-RCNN.

## Acknowledgements

We thank, Dr. Hidalgo and P. Lopez-Casas from Therapeutic Targets Laboratory at CNIO for providing us with panCAFs and Dr. de la Pompa and Edgell for providing MEVEC and EAHY926 cells, respectively. Dr. J. Pastor, and C. Blanco from Experimental Therapeutics Programme at CNIO for providing drugs, Dr. D. Filgueiras, R. Rouco and J.M. Alfonso-Almazán for help obtaining primary fibroblasts from pig hearts. We also thank Dr. J. Dorronsoro and A. Torres from machine learning group at UAM for helpful discussions on deep learning strategies. Editorial assistance was provided by Simon Bartlett.

## Supporting Information

**S1 Fig. Plate layout and CNN Nuc_Ring readout evaluation.** (A) Plate layout corresponding to a reference experiment used for both training and testing CNNs, where cells from indicated wells were used to create the training dataset with *healthy* (green) and *toxicity affected* (red) labelled cells coming from untreated wells and wells treated with the highest drug concentrations, respectively. (B) Plots display individual well toxicity readouts and the 5-Fluorouracil dose-response curve, including the EC_50_, from CNN_Nuc_Ring toxicity prediction. For each well, toxicity readouts were obtained by computing Z-scores (normalizing to DMSO-treated wells) with adjustment of the sign to display toxic effects as positive values. Z-scores > 3 represent toxic hits.

**S2 Fig. Evaluation of (R)CNN deep-learning toxicity-assessment approaches.** HL1 (A) and MEVEC (B) cells treated or not (-) with DMSO or the indicated concentrations of drugs (µM) were processed as described in the Materials and Methods. Representative images are shown of untreated cells. Plots display mean toxicity readouts of four replicate wells, obtained from the percentage of *healthy* cells predicted by the CNN_Nuc (Tox_CNN) or RCNN (Tox_RCNN) mixed models, and from nuclei counting by standard image segmentation (Num Nuc), or by RCNN-based automated detection (Num Nuc RCNN). For each well, toxicity readouts were obtained by computing Z-scores (normalizing to DMSO-treated wells) with adjustment of the sign to display toxic effects as positive values.

**S3 Fig. Confirmation of (R)CNN-predicted toxic hits.** Primary cardiac fibroblasts treated or not (-) with DMSO or the indicated concentrations of drugs (µM) were processed as described in the Materials and Methods. Boxplots of per-well toxicity assessments in culture wells from established measurements (A-C), and corresponding individual well toxicity readouts (D-F), obtained from Caspase 3/7 nucleus:cytoplasm ratio (Casp Nuc/Cyto) (A,D), Mitotracker cytoplasmic intensity (Mito) (B,E), and nuclei counting (Num Nuc)(C,F). Data are from 4 replicate wells of the same experiment. For each well, toxicity readouts (D-F) were obtained by computing Z-scores (normalizing to DMSO-treated wells) with adjustment of the sign to display toxic effects as positive values.

**S4 Fig. Validation of (R)CNN as drug toxicity screening tools.** Pancreatic CAFs treated with 60 compounds at the indicated concentrations (µM) were processed as described in the Materials and Methods. Plots display mean toxicity readouts of four replicate wells, obtained from the percentage of *healthy* cells predicted by the CNN (Tr_Tox_CNN) and RCNN (Tr_Tox_RCNN) mixed models after transfer learning, and from nuclei counting by standard image segmentation (Num Nuc), or by RCNN-based automated detection (Num Nuc Tr_RCNN). For each well, toxicity readouts were obtained by computing Z-scores (normalizing to DMSO-treated wells) with adjustment of the sign to display toxic effects as positive values.

